# Preventing *E. coli* biofilm formation with antimicrobial peptide surface coatings: recognizing the dependence on the bacterial binding mode using live-cell microscopy

**DOI:** 10.1101/2023.08.22.554337

**Authors:** Adam Hansson, Eskil André Karlsen, Wenche Stensen, John-Sigurd Svendsen, Mattias Berglin, Anders Lundgren

## Abstract

Antimicrobial peptides (AMPs) can kill bacteria by destabilizing their membranes, yet, to translate these molecules’ properties into a covalently attached coating is challenging. Standard microbiology methods do not work well for grafted AMPs, particularly it is difficult to distinguish the AMPs’ bactericidal potency from factors relating to bacteria’s binding behavior, e.g., which type of and how persistent bacteria-surface contacts that is necessary. Here we present a method combining live-cell microscopy and microfluidics to study the response of *E. coli* challenged by the same small AMP either in solution or grafted to the surface through click chemistry. The AMP coating initially suppressed bacterial growth as strongly as AMPs in solution. While AMPs in solution eventually killed the *E. coli* bacteria, those binding to the AMP coating changed contact mode one hour after binding and then became insensitive to it. The transition depended on binding-induced expression of Type 1 fimbriae, which limits contact between the AMPs and the *E. coli* outer membrane. By quantifying several different factors contributing to the antibacterial efficacy, these measurements provide a holistic understanding of how antibacterial surface coatings function. We therefore expect this tool to be important for the design of elaborate antibacterial coatings that can reduce the need for antibiotics and thus contribute to slower spreading of antibiotic resistance genes.

## Introduction

The rise of antibiotic resistance in common pathogens is a looming threat to patients worldwide as it disarm us our most potent tools against infectious diseases^1^. In medical settings, the issue is exacerbated by bacteria’s tendency to colonize and form resilient biofilm on abiotic surfaces found in medical devices such as implants, catheters and medical instruments^2^. Biomaterial-associated infections with resistant bacteria already pose a heavy burden on the resources of medical clinics and result in bad health outcomes for the patients^3–6^. To slow down the spreading of antibiotic resistance genes, the usage of antibiotics must decrease, a transition that would be easier if there were alternative ways to counteract biofilm formation.

One straightforward approach to escape biofilm formation is to inhibit bacterial colonization by chemically attaching substances that suppress bacterial growth, or binding, to the interface of biomaterials. It is less likely that bacteria will develop resistance to grafted substances, than substances leaking to the environment around the devices. Due to the covalent attachment, bacteria will experience a permanent, high concentration of antibacterial substance in the close vicinity of the surface. It will act selectively only on those bacteria that bind to the surface. Furthermore, in many applications the main aim will be to prevent the formation of resilient biofilm. Thus, also coatings that slow down growth of bacteria, or the development of biofilm-promoting phenotypes, selectively without killing may fulfil its purpose. Finally, the choice of antibacterial substance(s) may include molecules that have a very broad mode of action rather than targeting a specific biochemical mechanism. Although resistance development remains possible, it implies such large changes of the bacteria that it is ecologically disadvantageous in the long term.

Antimicrobial peptides (AMPs) is an archetype of the molecules that have potential to replace classical antibiotics in this type of applications^7–9^. These are amphiphilic peptides that can insert into the cell membrane(s) of bacteria, interact with the phospholipids and disturb membrane homeostasis^10^. Membrane binding is facilitated by a high content of cationic amino acids lending AMPs selective binding to bacterial membrane, which have more anionic lipids than other cell membranes^11^. Most AMPs can more efficiently inhibit Gram positive (G^+^) than Gram negative (G^-^) bacteria since the lipopolysaccharides (LPS) provide G^-^ bacteria an additional outer barrier that withholds the AMPs from entering the membrane^12^. When challenged by AMPs, to restore membrane function, general and specific envelop stress responses of the bacteria are activated^13^. Though, for high-enough exposure the bacteria cannot compensate for the damage and eventually die.

Most studied AMPs have a natural origin as being part of the innate defense of different procaryotic or eucaryotic species^11,14^. Natural peptides are commonly up to 50 amino acids in length and attain secondary structure such as α-helices or β-sheets for their active state^15^. Some different models have been suggested that describe how these AMPs disrupts the target cell membrane, e.g. through formation of transmembrane pores^16^. The use of natural AMPs for antibacterial products has been marginally successful; their length and dependence on native sequence makes their function sensitive to settings and enzymatic digestion, prone to cause off-target cytotoxicity, and lead to high production cost^9^. Through studies of the structure–activity relationship for natural AMPs, novel peptides that are smaller, more potent and more stable have been designed^17,18^. So-called small AMPs are made of a handful amino acids, mainly arginine (R) and tryptophane (W) providing the AMP hydrophobic and cationic properties, respectively^19^. The smallest pharmacophore was found to have only three residues^20^. Lacking apparent secondary structure, the antibacterial mechanisms of short AMPs remain elusive but is more likely due to interference with membrane protein function than formation of pores^21–23^.

The advent of small, stable and, relatively, cheap AMPs is a major step towards realization of AMP-based therapies and antibacterial products. Interestingly, the efficacy and susceptibility profiles of short AMPs is very sensitive to minor alteration of the order and frequency at which amino acids appear^19^ to peptide cyclization^24^ and to the position and stereochemistry of synthetic side-chains added to the pharmacophore^25^. In a recent study, different covalently surface-attached short AMPs could to varying degree suppress the colonization of G^+^ bacteria *Staphylococcus epidermidis*^26^. The efficacy of different coatings h however did not correlate to the AMPs’ MIC values in solution. It should indeed be recognized that the antibacterial efficacy of a coating relates to a combination of its ability to kill bacteria, to reduce surface growth, to prevent binding and to inhibit biofilm-specific phenotypes. MIC values obtained from small batch cultures might therefore not represent the potency of an antimicrobial peptide to prevent growth on surfaces and biofilms^27^. A good method to assist development of a coating should provide quantitative information about all modes of action, preferably under conditions that resemble that of the end-usage.

The current industrial standard test for antimicrobial surfaces (ISO 22196, ASTM E2149-13a, MBEC Assay®) is designed to measure released antimicrobial agents^28^. A method better suited for coatings of grafted AMPs is the CERTIKA test, which aim to measure the release of bacterial cells from a biofilm established on the test surface^29^. Most popular are fluorescence microscopy of surfaces immersed in bacterial solution for some time and then treated with so-called live/dead stain. These methods are commonly implemented as artificial end-point measurements without monitoring of the process dynamics. The outcome is therefore sensitive to the exact growth conditions, the way washing is done, and which type and for how long time staining is done^30–32^. Time-resolved live microscopy of single bacterial cells featuring fluorescent probes indicating membrane permeability, oxidative state etcetera have proven to efficiently detail the mechanisms of AMPs provided in solution^21,33,34^ and recently on antimicrobial polymer coatings^35^. The corresponding procedures and analysis are, however, complex and the throughput is low. Thus, there is an imminent need for medium-throughput methods that can both test the efficiency of novel AMP coatings and simultaneously deliver information about their mode of action to guide how they can be best used to create antibacterial materials.

Herein, we have built a microfluidic-based platform for live cell microscopy and subsequent automated image analysis using simplest possible microscopy without any labels or probes. As proof-of-concept, we synthesized an azide-conjugated version of the potent tripeptide AMP AMC-109^36,37^, which was tethered to the floor of the microfluidic channel using copper-catalyzed alkyne–azide cycloaddition (click reaction)^38^ and analyzed the binding to and growth of *E. coli* on this surface. To distinguish effects specific to details of the tethering from those specific to the AMP, bacteria were also bound to natural, mannose-modified surfaces and charged with the original pharmacophore and its azide-modified counterpart, respectively, in solution. This allowed us to extract qualitative and quantitative information on how an AMP coating works and its power to counteract biofilm. Importantly, bacterial responses beyond life or death could be distinguished, providing a holistic view of the coating’s antibacterial properties.

## Results

### Coating of microfluidic channels with AMPs through click reaction

AMC-109 (also known as LTX-109) is a synthetic tripeptide AMP that has proven efficient against common bacterial and fungal pathogens^36,39–41^. Its primary structure corresponds to R–W–R, giving the peptide a net charge of +3. The central tryptophane is modified with three bulky *tert*-butyl groups and the C-terminal is capped with an ethylphenyl group, which enhances the overall hydrophobicity of the peptide and protect it against enzymatic digestion^42,43^ (Figure 1a). To make the AMP surface coating of this pharmacophore, we created the peptide AMC-25-04 by attaching an azide-terminated short poly(ethylene) glycol (n=4, PEG) linker to the N-terminal of AMC-109, which also reduced its net charge to +2 (Figure 1b). The final structure was confirmed by NMR (Supplementary Figure 1) and mass spectrometry, and the product was purified to >95% using HPLC.

**Figure 1:**
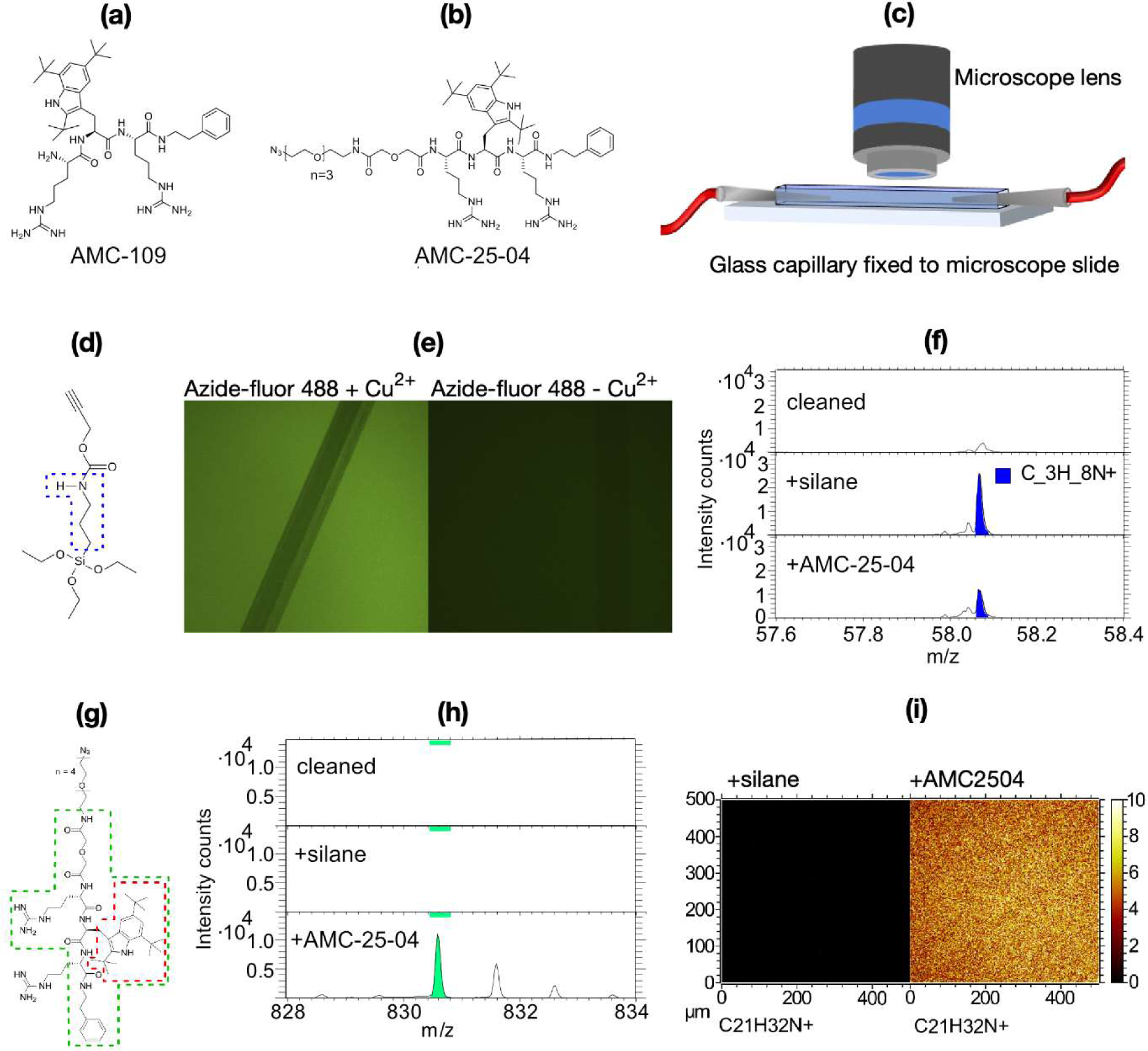
Surface modification of microfluidic channels. **(a)** Chemical structure of AMC-109. **(b)** Chemical structure of AMC-25-04. **(c)** Cartoon illustrating how the microfluidic channel was set up. **(d)** Chemical structure of O-(propargyloxy)-N-(triethoxysilylpropyl)urethane. The blue broken line indicates the potential origin of the fragment C_3_H_8_N^+^. **(e)** Fluorescence micrographs of silane-modified glass showing a glass surface after modification with Azide-fluor 488 in the presence of all reagents needed for click reaction (left panel), and a negative control where copper ions were omitted (right panel). The surfaces were gently scratched with a steel needle. **(f)** Part of ToF-SIMS spectrum showing total intensity counts of a clean glass surface (top), a surface modified with silane (center), and a surface modified first with silane and then with AMC-25-04 through click reaction (bottom). The blue-shaded peak corresponds to the ionized fragment C_3_H_8_N^+^ (m/z≈58.07, cf. panel **[d]**). **(g)** Chemical structure of AMC-25-04 where the ionized fragment with m/z≈830.6 is indicated by a green broken line and the fragment C_21_H_32_N^+^ (m/z≍298.8) corresponding to tri-tert-butyltryptophane is indicated by a red broken line. **(h)** Part of ToF-SIMS spectrum analogous to that of panel **(f)** highlighting the green-shaded peak corresponding to the fragment with m/z≈830.6 from the AMC-25-04 molecule. **(i)** ToF-SIMS micrographs showing the distribution of the ionized fragment C_21_H_32_N^+^ over an area of 0.5×0.5 mm^2^ (256×256 pixels) of surfaces modified with only silane (left) and silane+AMC-25-04 (right). The color scale shows the number of fragments detected for each pixel.

Bacterial binding and growth experiments were done using microfluidic channels of glass (Figure 1c). The inner walls of the capillary were modified with a silane presenting an alkyne functional group that allows orthogonal binding of azide-functional molecules through copper-catalyzed click reaction^38^ (Figure 1d). Fluorescence microscopy showed that azide-conjugated fluorophore readily bound to silane-modified surfaces in the presence of all reagents needed for click reaction, but very sparsely if copper ions were omitted (Figure 1e) confirming the reactivity of the silane modification.

Two types of molecules were attached to the glass surfaces using click chemistry. Azide-conjugated D-mannose was used for positive control experiments without AMPs, and for experiments where surface-bound bacteria were subject to AMPs in solution, respectively. Wild type *E. coli* can readily bind to D-mannose coated surfaces *via* their type-1 fimbriae^44^. AMP-coated surfaces were made by attachment of AMC-25-04. To verify the presence and homogeneity of the AMP coating we used time-of-flight secondary ion mass spectrometry (ToF-SIMS) analysis (Figure 1f-i). Glass surfaces modified with only silane showed high flux of an ionized molecular fragment C_3_H_8_N^+^ (Figure 1f). This corresponds to the central part of the silane (cf. Figure 1d). Upon binding of AMC-25-04, the flux decreased relative to the surface with only silane, which is expected due to shielding. Note that the peak visible in the spectrum for the clean surface corresponds to another, slightly heavier, fragment. Several unique fragments with high m/z values appeared after AMP binding. Figure 1h shows a spectrum highlighting an ionized fragment encompassing the full AMC-25-04 but for the PEG-linker and one of the arginine side chains (cf. Figure 1g). The fragment C_21_H_32_N^+^ corresponding to the artificial amino acid tri-*tert*-butyltryptophane (cf. Figure 1g) was found to coat the surface homogenously on the scale corresponding to that observable with optical microscopy (Figure 1i).

### Bacterial binding and growth on surfaces under favorable conditions

Bacteria dispersed in growth media were injected into the microfluidic channels for 20 minutes at a low flow rate, which facilitates their binding to the channel bottom. Bacteria solution was then replaced with pure growth media and the flow was increased to get a sufficient flux of nutrients to support fast growth of bound bacteria (cf. Method section & Supplementary Figure 2). The binding and subsequent growth was filmed for >3 hours at a rate of 2 fps with a microscope operated in brightfield mode. Automated image analysis was implemented using an in-house developed MATLAB program: the bacterial objects were distinguished by their shape and trajectories constructed by stitching objects in subsequent frames together. We chose to determine the bacterial length as our main characteristic since this is straightforward for rod shaped bacteria and its value is relatively unsensitive to noise or focus drift. Furthermore, the length and its derivate with respect to time, i.e. the growth rate, are good indicators of the metabolism of a rod-shaped bacterium and whether it is subject to stress^45–47^.

Figure 2 summarizes the analysis of a control experiment showing normal growth behavior of *E. coli* bound to a mannose-coated surface. In the first phase (<30 min) the number of bacteria increases due to binding, in the second phase (30–70 min) the number remains almost constant, and in the last phase (>70 min) the number increases exponentially as bacteria divide (Figure 2a). Figure 2b–c show results of the tracking procedure applied on the same bacteria as in Figure 2a. As detailed in the Method section, only traces >10 minutes were used to construct these plots. A new trace started if a bacterium appeared on a position not occupied in previous frames, or when the cell divided (Figure 2b). After binding, the bacteria remained slightly mobile moving in the direction of the flow but eventually binding strength increased, which is discussed further below. Figure 2c shows the data points of trajectories (color-coded) included in the analysis, plotted in the time domain. Few division events are seen before 70–80 minutes, later cells divide approximately every 30 minutes. Cells grow linearly in the beginning of a cell cycle and faster, almost exponentially, when approaching division, in line with previous observations^48^.

**Figure 2:**
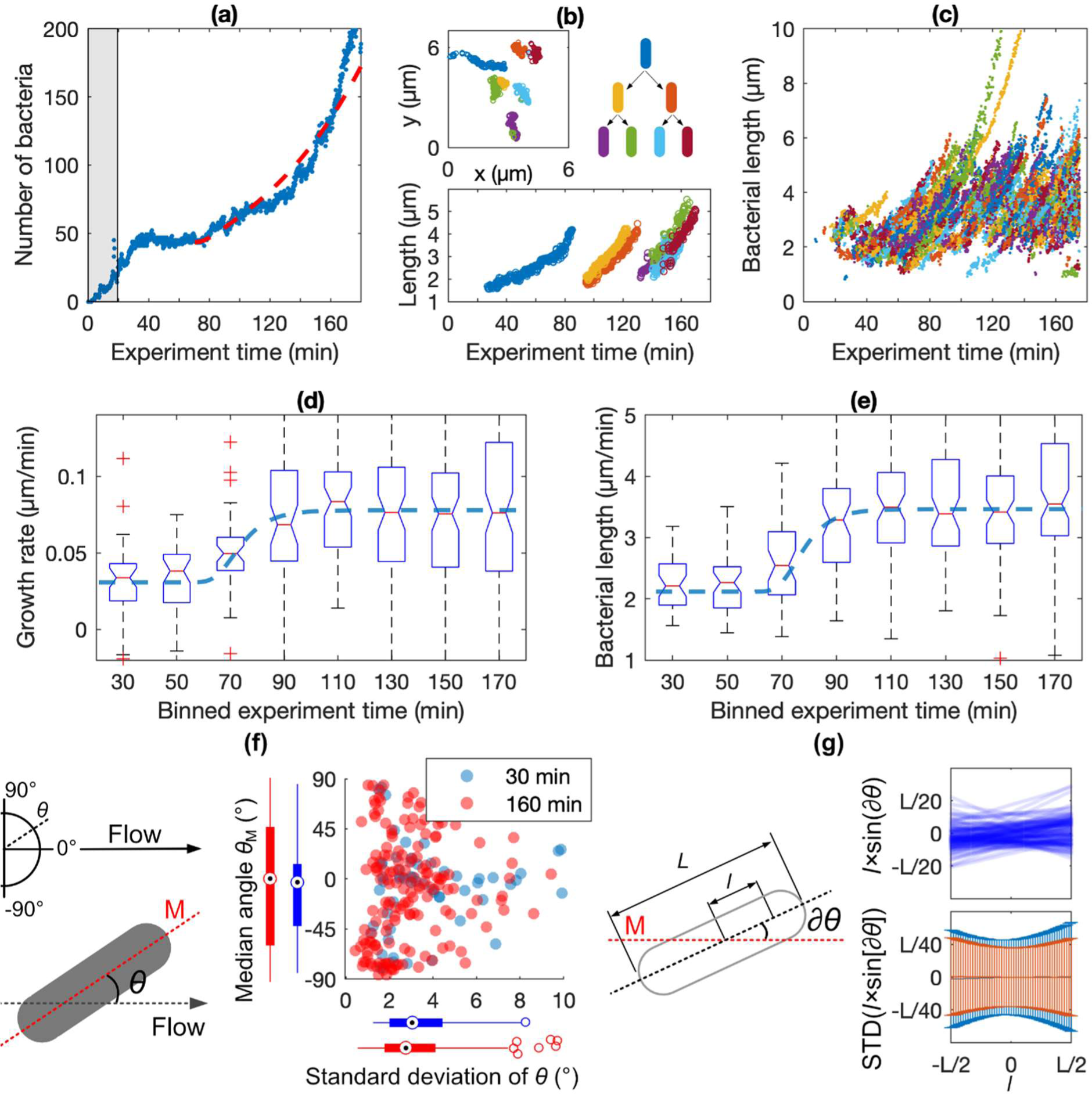
Method for analysis of binding and growth. **(a)** Number of identified bacteria on the imaged surface from the beginning to the end of the experiment. The gray-shaded region indicates the period of bacteria injection. The red broken line indicates the best fit of eq. 1 to the number data. **(b)** Selection of traces showing the formation of a microcolony plotted in the spatial (upper panel) and time (lower panel) domains, respectively. The mother-daughter relations are shown by color code. **(c)** Plot showing all data points, color-coded according to the tracing procedure, for which growth rates (GR) were calculated. **(d)** Boxplot showing the distribution of GRs of the bacteria present divided into 20-minutes intervals. The x-axis values indicate the center of each interval, e.g., 50 min indicates the time interval 40–60 minutes. **(e)** Boxplot showing the distribution of the lengths of the bacteria for which GR was presented in (d). **(f)** Cartoon and plot detailing the analysis of bacterial alignment, i.e., the angle θ between a bacterium’s major axis and the flow direction. The combined scatter and boxplot show the median angle, θ_M_, versus the standard deviation (STD) of θ for each bacterium at an early (30 min, blue points) and a late (160 min, red points) timepoint of the experiment. **(g)** Cartoon and plots detailing the analysis of bacteria’s wiggling movements around their median major axes. The scatter plot in the upper panel shows for a single example bacterium the instantaneous separations, l×sin(∂θ), between each position l along the bacterium’s major axis L and the median major axis M. The bar plot in the lower panel shows the distribution (standard deviations) of the instantaneous separations, l×sin(∂θ), for all positions l of all bacteria present early (30 min, blue bars) and late (160 min, red bars) in the experiment.

Growth rates (GR) were calculated from the gradient of the length versus time plots, and the average values obtained for each bacterium were binned as shown in Figure 2d. The average lengths of the same bacteria at the same time intervals are shown in Figure 2e. The first 40 minutes after binding, bacteria are small and grow slowly and the distribution of both GR and cell length is narrow. This behavior is a well-known response to surface adhesion, which make bacteria to collectively enter a lag-phase during which expression of treats related to planktonic life is downregulated and treats important for biofilm formation is upregulated^49^. After about 40 minutes the GR is accelerated and is eventually saturated at about twice its initial value. The cell length simultaneously increases and saturates at about 1.5 times its initial value. The distribution of GR and length is also broadened as cells grow at different rates during different phases of their cell cycle.

The number of bacteria on the surface will increase exponentially with time, with a rate constant depending on the duplication time, *T_2_*, and on the rate of bacteria release from the surface indicated by the mean binding time, *T_½_*. Since *T_2_* can be determined from the bacterial GR and length at division, *T_½_* may be estimated from a fit to the plot of the number of bacteria versus time (Figure 2a, red broken line), providing *T_½_*≈60 minutes for the experiment above (c.f. Method section). This approach is robust but approximative, since it does not take into account that binding strength may change with time. Hence, we also analyzed the interplay between the surface adhesion mode of the individual bacteria and the flow forces in the beginning (30 min) and at the end (160 min) of the experiment.

Just after binding, the rod-shaped bacteria tend to align their major axis approximately with the direction of the flow. With time, more bacteria attain a position approximately normal to the flow direction (Figure 2f). Bacteria attached with a large angle relative to the flow also tend to wiggle less than those aligning with the flow, as shown by the standard deviation of the binding angle. The realignment indicates that additional bonds, distributed along the bacterial rod, form between a bacterium and the surface. These bonds are strong enough to withstand the flow force that causes bacteria to align with flow. Analyzing the amplitude of the small motions around the median major axis of the bacterium can indicate the location and character of the attachment points under its body (Figure 2g)^50^. The *WT E. coli* binding to mannose-coated surfaces appear to rock slowly, rather than twist, symmetrically around their centers (Figure 2g [upper panel] & Supplementary Movie 1). The central part of the bacteria moves slightly less than the poles and the difference become even smaller towards the end of the experiment (Figure 2g, lower panel).

We suggest that the observed realignment of the bacteria away from the flow direction and the symmetric rocking motion are characteristic for fimbriae-mediated binding to the surface since fimbriae are relatively long and distributed evenly over the bacterial body. This hypothesis explains that binding is both flexible *and* homogenous along the bacterial rod. Likely, the adhesion pattern changes with time due to increased fimbriae expression after binding. This interpretation is supported by a similar analysis made on *E. coli* that overexpresses fimbriae, *E. coli-Fim+*. In that case, the initial adhesion pattern reassembles the final pattern observed for *WT E. coli,* and less change occurred over time (Supplementary Figure 3).

### Bacterial response to AMPs provided in the growth media

The normal growth of WT *E. coli* on mannose-coated surfaces was contrasted by very different behavior when the growth media applied after binding was supplemented with 100 µM (>MIC) of the AMPs AMC-109 or AMC-25-04. The bacteria subjected to AMPs immediately showed attenuated GRs and, finally, growth arrest (Figure 3a). AMC-25-04 lowers the initial GR by 24% and AMC-109 by 57% relative to the control (Figure 3b). The variation of the initial median GRs between replicates is small, particularly for the control and experiments with AMC-25-04. This is notable since *E. coli* cultures had reached different OD in the LB media before binding to the surface in the different replicates and thus likely also had different GRs (c.f. Method section). This shows that the attenuated GR values measured after binding only reflect the surface-adhered state of the bacteria and the AMP-containing environment in combination, not the previous growth conditions. The distribution of initial GRs of individual bacteria in a single experiment with AMPs is also narrow (Supplementary Figure 4). The GR attenuation is accordingly not due to some bacteria being more sensitive to the AMP and therefore reaching GR arrest early. The distribution of the individual GRs however broadens as the median GR starts to decline, indicating that the transition to GR arrest is abrupt on the single-cell level.

**Figure 3:**
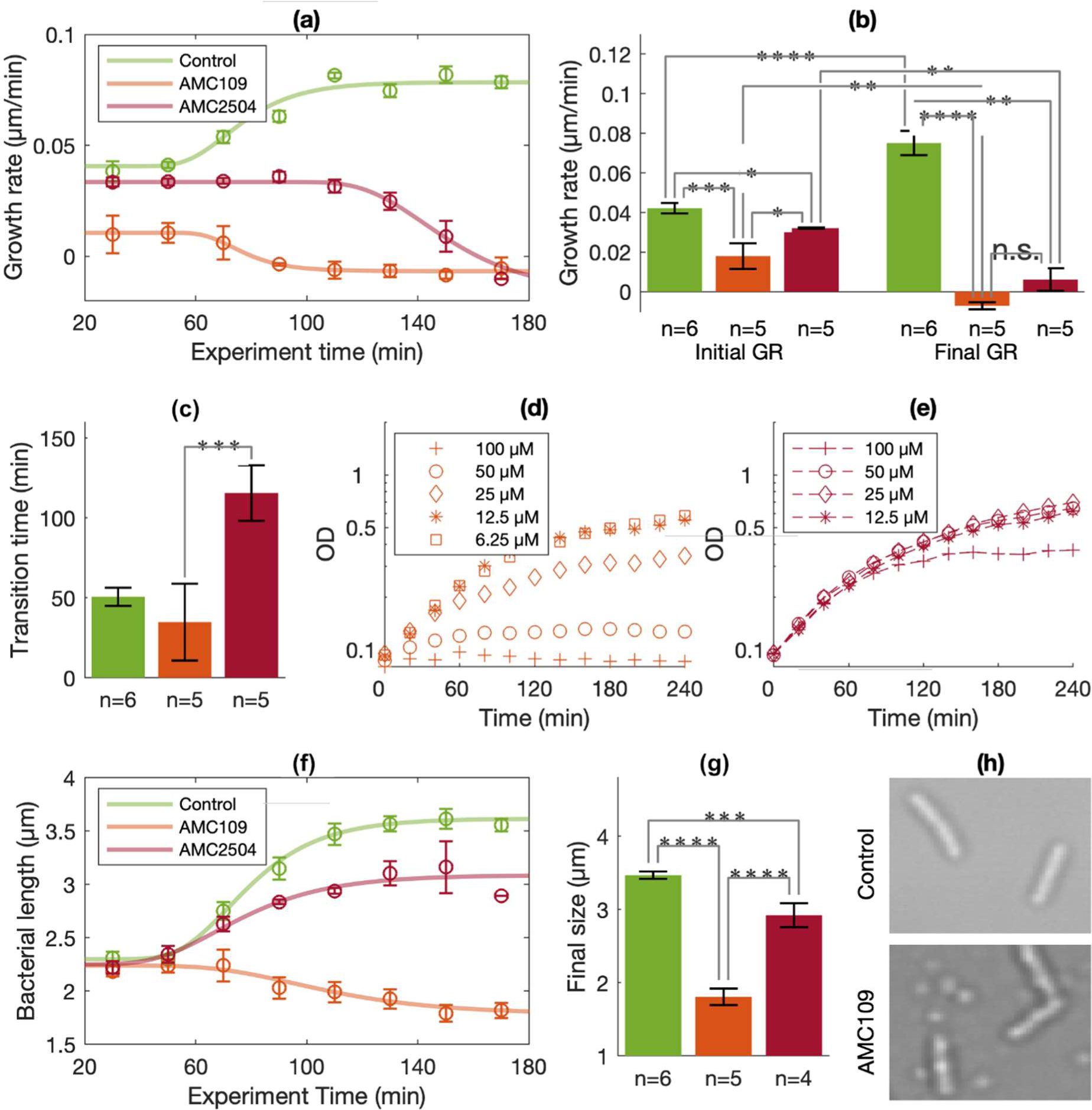
Efficacy of AMPs delivered in growth media. In all plots green color denotes control experiments (N=6), orange color denotes experiments with added AMC-109 (N=5), and red color denotes experiments with added AMC-25-04 (N=5). Statistical significance was tested by Student’s T-test where n.s. denotes not significant, * p<0.05, ** p<0.01, *** p<0.005 and **** p<0.001. **(a)** The plot shows the average and standard error (SE) of the median GR values for each bin in the boxplots (c.f. Figure 2d) of all individual experiments. **(b)** The bars show the average and SE of median GRs measured before (“Initial GR”) and after (“Final GR”) the transition time was reached in the individual experiment. **(c)** Bars show the average and SE of the transition times determined in each experiment. The transition times correspond to the inflection points of the Gompertz’s fits to the data in the boxplots (c.f. Figure 2d). **(d)** The plot shows batch-culture growth curves for E. coli in the presence of different concentrations of AMC-109. **(e)** The plot shows batch-culture growth curves for E. coli in the presence of different concentrations of AMC-25-04. **(f)** The plot shows the average and SE of the median bacterial size values for each bin in the boxplots (c.f. Figure 2e) of all individual experiments. **(g)** Bars show the average and SE of median sizes measured after the transition time was reached in the individual experiment. **(h)** Micrographs detail the appearance of E. coli towards the end of a control experiment (upper panel) and an experiment where AMC-109 was added (lower panel). Both bacteria and AMP nanoparticles (small round features) are visible in the lower micrograph.

The time elapsed from the addition of AMPs to growth arrest is 116 min for AMC-25-04 and 35 min for AMC-109 (Figure 3c). Taking either the initial GR attenuation, or the time to GR arrest as measures of antibacterial efficacy, both observations indicate that AMC-109 is about 3 times more potent than AMC-25-04. This result compares well with data obtained from batch-culture experiments (Figure 3d–e). Growth curves measured for *E. coli* charged with 100 µM AMC-25-04 and 25 µM AMC-109 showed similar progression. The time to GR arrest is also similar in live-microscopy and batch-culture experiments. Yet, the initial attenuation of GR relative the control observed for surface-bound bacteria subject to AMC-25-04 is not detectable in the corresponding batch culture experiments.

The progression of cell length was strongly impacted by the AMPs (Figure 3g–f). Most strikingly, AMC-109 makes cells shrink. This is partly due to bacteria dividing asymmetrically creating small cells, and partly due to an actual, abrupt cell shrinkage taking place as bacteria die (Supplementary Movie 2). These processes also lend bacteria irregular shape characterized by an uneven intensity distribution and less well-defined edges than normally growing *E. coli* (Figure 3h). The effect of AMC-25-04 was initially less drastic, causing the bacteria to grow smaller than in the controls. Decreased cell length homeostasis is indeed a characteristic of *E. coli* growing while subject to stress^47^. Towards the end of the experiment, similar behavior was observed for AMC-109. A surprising observation was the formation of AMP nanoparticles, appearing on the surfaces early in experiments with AMC-109 and later AMC-25-04 (Figure 3h, lower panel). These particles were also present in water-based solutions not containing bacteria, excluding that they are debris of dead cells. It was recently reported that AMC-109 can form 5 nanometer aggregate structures that potentiates its antibacterial action^23^. The particles observed here are much larger (>100 nanometer in diameter) and it is unclear whether they impact the efficacy of the AMPs.

### Bacterial response to AMP-coated surfaces

The *WT E. coli* bound readily to surfaces coated with AMC-25-04. Right after binding, the initial GR is attenuated by 26% relative to the GR of WT *E. coli* growing on mannose-coated surfaces (Figure 4a–b). The GR, however, increased steeply in a similar way and after the same lag time as seen for the controls. After 100 minutes, the GR of WT *E. coli* on AMP-coated surfaces catch-up with that measured in the control experiments; the final GRs are not significantly different. Interestingly, *E. coli* growing on the AMP-coated surfaces changed adhesion mode drastically during the experiment (Figure 4c). Initially, bacteria align strongly with the flow and wiggle much more than on the mannose-coated control surfaces. At the end of the experiment, however, the alignment and wiggling behavior is on a par with that seen for fimbriae-mediated binding to mannose (cf. Figure 2).

**Figure 4:**
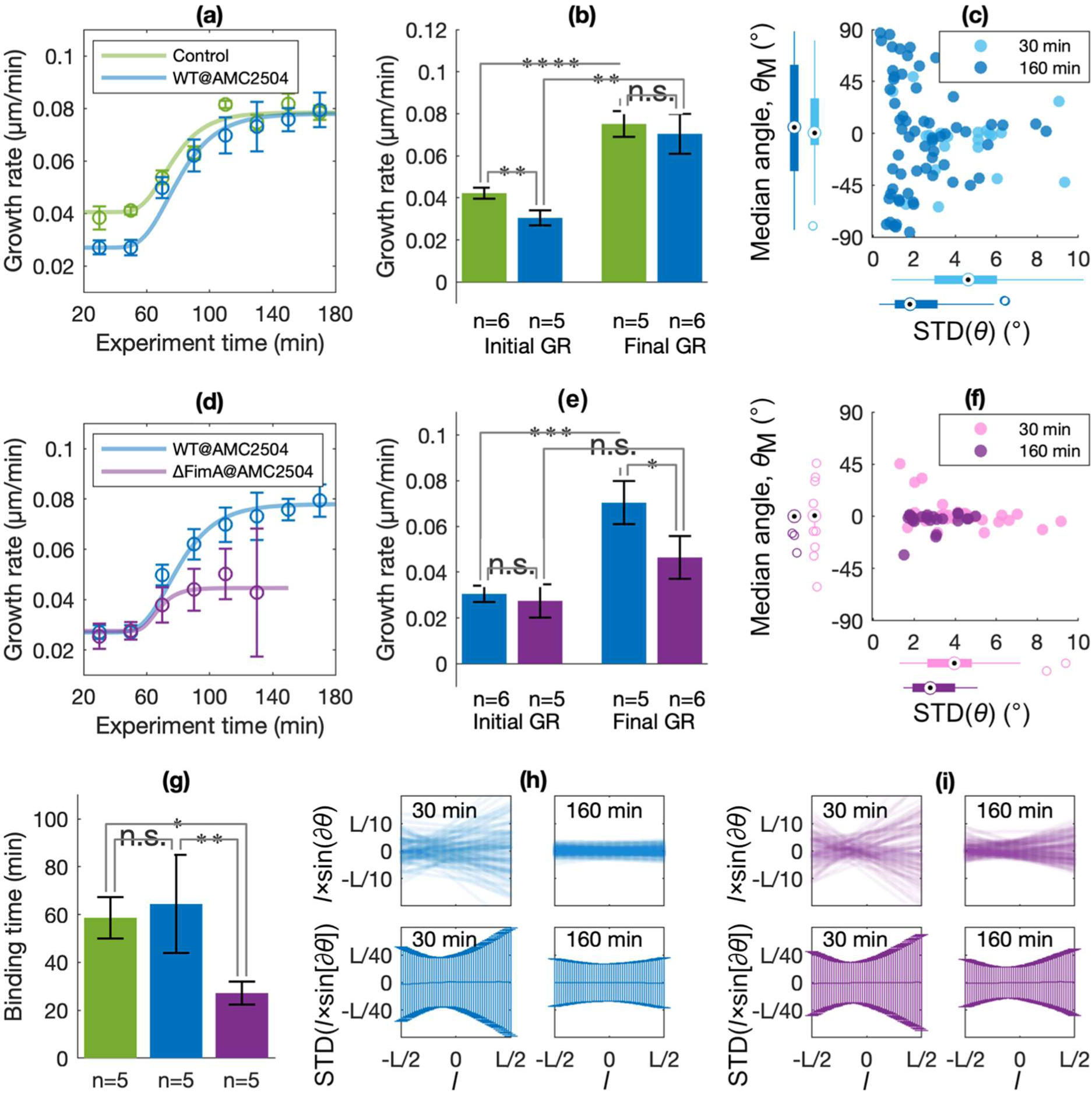
Efficacy of AMP-coated surfaces. In all plots green color denotes control experiments (N=6), blue color denotes experiments with WT E. coli (N=5), and violet color denotes experiments with fimbriae-deficient E. coli-ΔFimA (N=5). Statistical significance was tested by Student’s T-test where n.s. denotes not significant, * p<0.05, ** p<0.01, *** p<0.005 and **** p<0.001. **(a)** The plot shows the average and standard error (SE) of the median GR values for each bin in the boxplots (c.f. Figure 2d) of all individual experiments. **(b)** The bars show the average and SE of median GRs measured before (“Initial GR”) and after (“Final GR”) the transition time was reached in the individual experiment. **(c)** The combined scatter and boxplot shows the median angle, θ_M_, versus the standard deviation (STD) of θ for all bacteria at an early (30 min, light blue) and a late (160 min, dark blue) timepoint of the experiment (c.f. Figure 2f). **(d)** The plot shows the average and standard error (SE) of the median GR values for each bin in the boxplots (c.f. Figure 2d) of all individual experiments. **(e)** The bars show the average and SE of median GRs measured before (“Initial GR”) and after (“Final GR”) the transition time was reached in the individual experiment. **(f)** The combined scatter and boxplot show the median angle, θ_M_, versus the standard deviation (STD) of θ for all bacteria at an early (30 min, pink) and a late (160 min, violet) timepoint of the experiment (c.f. Figure 2f). **(g)** Bars show the average and SE of bacteria’s mean binding times, T_1/2_, determined in the different experiments. **(h&i)** Scatter plots in the upper panels show for a single example bacterium the instantaneous separations, l×sin(∂θ), between each position l along the bacterium’s major axis L and the median major axis M. The bar plots in the lower panels show the distribution (standard deviations) of the instantaneous separations, l×sin(∂θ), for all positions l of all bacteria (c.f. Figure 2g).

The covalently attached AMPs extend just a few nanometers from the surface. The diminishing efficacy of the AMP-coating observed after the time-lag may thus be explained by the shifting adhesion mode. If numerous, micrometer-long fimbriae protrude from the bacterial body it will likely limit the access of the AMPs to the bacterial membrane. To test if increased production of Type 1 fimbriae post-binding impacts bacteria’s sensitivity to the AMP-coating, we studied the surface growth of a mutant, *E. coli-ΔFimA*, that do not produce fimbriae (Figure 4d-e). Initially, *E. coli-ΔFimA* grow at the same, reduced rate as WT *E. coli* on the AMP-coated surfaces. However, in contrast to WT *E. coli,* the GR does not increase to the same extent after the lag-phase for the bacteria without Type 1 fimbriae: the final GR was 34% lower than that measured for WT *E. coli*.

The cell alignment analysis showed that non-fimbriated *E. coli* aligned strongly with the flow (Figure 4f), similarly to the WT *E. coli* (c.f. Figure 4c). In contrast, no realignment was observed for *E. coli-ΔFimA*, and wiggling decreased less than for WT *E. coli* towards the end of the experiment. Notably, although *E. coli-ΔFimA* bound readily to the AMP-coated surface, many bacteria left upon division, in particular the daughter cells. The characteristic binding time, T_1/2_, is only 27 minutes, which is less than half of that seen for WT *E. coli* (Figure 4g). This is also less than the final duplication time, T_2_, measured for *E. coli-ΔFimA* on the surfaces (67 minutes) and thus the total number of bacteria decreases over time. Consequently, after 140 minutes reliable analysis of GR was not possible (Figure 4d).

The results obtained for WT *E. coli* and *E. coli-ΔFimA* growing on the AMP coating indicate that the bacterial phenotype that align with the flow and wiggle a lot is affected by the AMP coating, resulting in a reduced GR. This was observed for WT *E. coli* early in the experiment, and always for *E. coli-ΔFimA*. Another feature of this phenotype is that bacteria’s small movements can be described as twisting around a fixed point located somewhat upstream the bacteria’s center points, as seen in the plots for WT *E. coli* in the beginning of the experiment (Figure 4h [left panels], Supplementary Movie 3) and for *E. coli-ΔFimA* both in the beginning and in the end of the experiment (Figure 4f, Supplementary Movies 4-5). This pattern of movement is characteristic for an *E. coli* bacterium adhering to a surface via a patch located on its body^50^, i.e. in the proximity of the outer membrane. In the above experiments, only those bacteria that express Type 1 fimbriae realign away from the flow direction during growth. This behavior was observed for WT *E. coli* towards the end of the experiment. The corresponding analysis of the bacteria’s small motions indeed revealed a pattern like that observed for WT *E. coli* binding to mannose-modified surfaces (c.f. Figure 2g) characterized by a slowly symmetrically shaking motion (Figure 4h [right panels], Supplementary Movie 6). This implies that WT *E. coli* initially bind to the AMP-coated surfaces through adhesion patches on their bodies, but gradually shift binding mode to bind via Type 1 fimbriae distributed evenly along their bodies.

## Discussion

The combination of live-microscopy and microfluidics used in this work is superior to classical batch-culture methods in measuring the efficacy of antibacterial surface coatings: We could distinguish and quantify the contribution of several different antibacterial mechanisms under settings that reasonably well mimic the intended final implementation. Using microfluidics also removes inherent problems associated with many classical methods such as the inoculum effect^51^ and variability caused by manual interference (e.g. rinsing and staining steps) during the experiment. Thus, the obtained data will be of high quality making it possible to detect also small differences between different coatings. This is important, not at least to enable a rational design strategy. Furthermore, our implementation features simplest possible microscopy, automated analysis, and generally applicable click-chemistry for surface modification, making this platform suitable for upscaling.

In all experiments featuring AMPs the initial growth rate (GR) was significantly lower than in the control experiments. It can be argued, from different perspectives, that the initial GR on surfaces indeed is a good measure of an AMP’s potency. For the two AMPs tested in our experiments, the initial GR correlated with the time needed to reach complete growth arrest, yet more AMPs must be analyzed to establish if this relationship holds generally. From a measurement perspective, it is interesting that upon surface binding the bacteria “reset” to prepare for biofilm formation. This gives rise to narrow distributions of GRs during this phase of the experiments, both between individual bacteria and between different experimental replicates, providing a stable baseline for measuring deviations of GR caused by AMPs. It is not well-understood how the bacteria sense that they are bound to a surface, but adhesion lead to activation of Cpx- and σ^E^-regulated envelop stress responses (ESR) of *E. coli* bacteria^49,52^. The same ESR systems provide the bacteria’s first response to AMPs^53^. Potentially, attenuation of GR due to the summed impact of surface binding and AMPs is more easily measured than the effect of AMPs alone, at least if the latter is comparably small. The initial GR can also be measured relatively fast, reliable values can be acquired within one hour. Finally, the GR after surface binding is also a particularly relevant measure if the final aim is to establish AMPs for the purpose of antibacterial coatings since it directly reflects the biofilm formation process.

The *E. coli* bacteria are seemingly less sensitive to surfaces coated with small AMPs than *S. epidermidis,* which was investigated in a previous study using the CERTIKA method^26^. At first sight, this result is not surprising since G^-^ bacteria are generally less susceptible to AMPs than G^+^ bacteria due to the LPS of G^-^ bacteria obstructing the AMPs from inserting into the membrane core. It is unclear if this mechanism holds also for a covalently attached AMP coating: While the uptake of AMPs from bulk media is rate-limited by the lipid composition of the bacteria’s membrane^23^, the amount of AMPs protruding into the membrane at the interface between bacterium and coating depends also on the grafting density of AMPs, the geometry of the interface, and the adhesion forces that pull the bacterial membrane and substrate together. Judged by their GRs, *E. coli* were initially equally affected if AMC-25-04 was provided in a high concentration in the growth media, or if it was presented as surface coating. As shown herein, only a minor part of the membrane, the adhesion patch, is in contact with the surface coating. Accordingly, bacteria appear equally stressed when their entire membrane is exposed to the AMP in solution as when just a minor fraction of it is in contact with the AMP coating. Due to the covalent grafting, the AMP concentration may be locally very high and persistent in the contact patch. It can thus be speculated that the cells better tolerate (per membrane area unit) the AMPs absorbed spontaneously from bulk than AMPs pushed into the membrane and, consequently, the overall impact on the GR is similar. From a materials design perspective this finding is interesting since it shows that an antibacterial coating can function without a large contact area between the bacterial body and the material.

Another factor that clearly contributes to that *E. coli* bacteria show tolerance towards the AMP coating is their ability to change binding mode during biofilm adaptation. The planktonic phenotype, which still dominates initially after binding, is sensitive to the AMP coating while the phenotype evolving during the post-binding lag-phase, is not. The difference relates to a transition from an adhesion mode where a patch of the bacterial body is in contact with the surface, to a mode where the bacteria bind to several distributed contact points via fimbriae. Notably, fimbriae are comparably stiff structures and if present they will push a bacterium slightly away from the surface^54^. This will lower the chance that the bacterial outer membrane gets in contact with the coating. Type 1 fimbriae are the most common type of pili on *Escherichia*, but it is also present on other G^-^ bacteria, e.g., *Klebsiella* and *Pseudomonas*, implicated in biofilm formation on biomaterials. Many other types of pili found on G^-^ have similarly stiff structures^55^ and may thus give rise to the same effect. The mechanical properties of pili of G^+^ bacteria have not been studied to the same extent. This result emphasizes that bacteria is most vulnerable during a time-window right after binding before development of the biofilm phenotype occurs. It also shows that a deeper understanding of the details of the bacteria–substrate interface is needed so materials and grafting strategies adapted to bacteria’s binding mode can be developed. Different responses to surface binding may ultimately form the basis for the design of coatings with selective antibacterial properties.

## Methods

### Synthesis of AMC-25-04

Amino-azido derivate of PEG200 was prepared as described by Jiang *et al.*^56^ and reacted with diglycolic anhydride to provide azido-PEG-COOH linker. One equivalent (EQUIV) azido-PEG-COOH was coupled to 1 EQUIV AMC-109^42^ (Amicoat A/S, Norway) using 1.2 EQUIV HBTU and 4.8 EQUIV TEA to gain AMC-25-04. The crude peptide was purified by preparative HPLC and lyophilized to yield TFA-salt. The purity was >95% as determined by HPLC. The product was confirmed with NMR (Supplementary Figure 1) and ESI-MS: (MH^+^, monodisperse, calculated for C_55_H_90_N_15_O_9_: 1104.705, observed 1104.7009).

### Microfluidic channel assembly and silanization

Microfluidic channels were prepared by mounting a square glass capillary with ID 0.8×0.8 mm^2^ (VitroCom, USA) on a microcopy slide using UV-curable adhesive (NOA 68, Norland Products inc., USA). The capillary was connected to tubing at both ends using Gel-loading Pipette Round Tips (VWR). The channels (and coverslip glass surfaces used for fluorescence microscopy and TOF-SIMS analysis) were cleaned by immersion overnight with Hellmanex III cleaning solution (2%, Hellma GmbH, Germany) followed by immersion in Sulfuric acid (2M, Sigma-Aldrich 99.9%) for one hour, followed by extensive rinsing with water (Milli-Q, Merck Life Science). The cleaned glass was rinsed with ethanol (99.5%, Solveco, Sweden) and then silanized by immersion in 10% solution of O-(Propargyloxy)-N-(triethoxysilylpropyl) urethane (90%, ABCR, Germany) in ethanol (99.5%, Solveco, Sweden) for one hour.

### Click-chemistry modifications of glass surfaces

The silanized capillaries and coverslips were modified with AMC-2504 or α-mannose-PEG3-azide(>95%, Sigma-Aldrich) or Azide-fluor 488 (>90%, Sigma-Aldrich) using copper-catalyzed alkyne-azide cycloaddition (CuAAc, click-chemistry)^38^. The glass modified with the alkynated silane was conjugated to the different azidated molecules by immersion for 10 minutes in click reaction solution containing 33 µM azidated reactant, 17 mM Guanidine hydrochloride (Sigma-Aldrich), 75 µM CuSO_4_ (Sigma-Aldrich), 250 µM Tris(3-hydroxypropyltriazolylmethyl)amine (THPTA, Tokyo Chemical Industry Co., Ltd.) and 500 µM Ascorbic acid (Merk) diluted in PBS buffer (pH 7.4).

### Verification of surface modifications

The glass surface modifications were verified with fluorescence microscopy and Time-of-Flight Secondary Ion Mass Spectrometry (ToF-SIMS). Coverslip glass modified with silane was modified with Azide-fluor 488 using click reaction solution with and without added CuSO_4_ (see above). After rinsing with water, the surfaces were scratched with a needle to provide some contrast. Surfaces were imaged immersed in water using an upright fluorescence microscope (Axioskop 20, Carl Zeiss Microscopy) equipped with a water-immersion objective (W plan-apochromat 63x/1.0, Carl Zeiss Microscopy) using the same exposure time. ToF-SIMS (TOFSIMS IV, IONToF GmbH, Germany) was used to analyse the presence and spatial distribution of peptide fragments before and after their surface attachment to the silanized glass. The instrument was operated using 25 keV Bi^3+^ primary ions at a pulsed current of 0.1 pA (cycle time 150 µs, width 1.2 ns). The samples were analyzed in the bunched mode at an analysis area of 500 × 500 µm^2^ (resolution 256 pixel). The average of 25 scans at an acquisition time of 100 sec was used when acquiring data using SurfaceLab version 6.7 software from IONToF GmbH (Münster, Germany).

### Bacteria and growth media

Versions of the *E. coli* wild-type (WT) strain MG1655 were used throughout all experiments. *WT E. coli* were provided green fluorescence (GFP) and resistance to Kanamycin by transformation with plasmid pBE1-mGPFmut2^57^. *E. coli-Fim+* that overexpress Type 1 fimbria was made by transformation with plasmid pPKL91, which promote fimbriae expression by increasing the intracellular concentration of the regulating protein FimB^58^. *E. coli-ΔFimA* that lack Type 1 fimbriae was made by deletion of the *fimA* gene encoding the major structural protein (FimA) of the fimbria from the chromosome of *E. coli* MG1655. For this, *ΔfimA::kan* was transduced from the KEIO collection strain JW4277^59^ by standard methods to MG1655, selected for Kanamycin resistance, and the locus verified by PCR. The fimbriated and non-fimbriated phenotypes were confirmed by yeast agglutination. The bacteria were kept in deep-frozen glycerol stocks and on weekly basis plated and grown on LB-Agar. For live-cell microscopy experiments, a single colony was selected from a plate, inoculated into LB media supplemented with 50 µg/ml Kanamycin and grown overnight at 37°C. The overnight cultures were gently centrifuged to remove aggregated bacteria. The supernatant, which typically had OD_600_≈0.1, was diluted 1:1 in fresh LB media supplemented with Kanamycin and grown at 37°C until 0.35< OD_600_<0.6.

### Live-cell microscopy

The microfluidic channel was mounted under a microscope (Axioskop 20, Carl Zeiss Microscopy) equipped with a stage heated to 37°(SKE, Italy). The channel was connected to a syringe pump (NE-300, New Era Pump Systems) via polypropylene tubing. A pipette tip attached to the tubing by UV-curable adhesive (NOA 68, Norland Products inc., USA) worked as adapter between the tubing and the pipette tip attached to the channel. The lower surface of the channel was imaged using a water-immersion objective (W plan-apochromat 63x/1.0, Carl Zeiss Microscopy) and footage was acquired using a microscope camera (Axiocam 305 Colour, Carl Zeiss Microscopy) at 2 fps. Bacteria in LB media (see above) was taken directly from the incubator, transferred to a syringe, and injected into the channel, first 10 minutes at a flow rate of 100 µl/min to equilibrate the system, and then for another 10 minutes at a lower flow rate of 20 µl/min to promote bacterial binding. The syringe was then replaced with a new syringe containing LB media, or LB media supplemented with 100 µM AMPs, which was injected at flow rate 100 µl/min for approximately 3 hours. Notably, with the present channel dimensions, the flow rate of 100 µl/min translates into a flow speed of approximately 30 µm/s at 1 µm separation from the surface where the bacteria sit^60^ (Supplementary Figure 2). At lower flow speed *E. coli* grow significantly slower and do not form microcolonies as (data not shown). If channels with other dimensions are used it is therefore important to recalculate and adapt the flow rate accordingly.

### Image analysis and segmentation

Image analysis of footage was done with an automated workflow written in Matlab (MATLAB Version: 9.13.0.2049777 [R2022b], The MathWorks Inc., USA) featuring functions of the Image Processing Toolbox. For analysis of GR, non-bound bacteria were excluded from analysis by averaging every 20 consecutive frames, reducing the effective frame rate from 2 fps to 0.1 fps. For analysis of bacteria’s small motions around their major axis, footage was analyzed at the original frame rate for periods of 2 minutes.

Images were corrected for uneven illumination and background was subtracted. The resulting image was used to construct a mask by, in the following order, subtraction of a user-set constant value, two rounds of median filtering, marker-controlled watershed segmentation, and application of shape/size criteria. Using this mask, properties of the individual bacteria were extracted; the length was measured as the major axis of the best-fitting centroid. Bacteria in subsequent frames were stitched together to trajectories if their footprints overlapped and their length changed <25%. To handle issues with bacteria temporarily “disappearing” from the segmented mask, e.g., due to focus drift, a bacterium remained member of the same trajectory although it was not visible for some short time if it reappeared with unaltered appearance. Else, new trajectories start if bacteria divide or appear on new positions.

### Analysis of growth rate and mean binding time

The evolution of population growth rate, *GR(t)*, cell length, *L(t)*, and the overall mean binding time, *T_½_*, were extracted from trajectories of growing cells using Matlab (MATLAB Version: 9.13.0.2049777 [R2022b], The MathWorks Inc., USA) featuring functions of the Curve Fitting Toolbox and the Statistics and Machine Learning Toolbox. The momentaneous GR of a bacterium was determined by the best linear fit to cell length data extending +/- 5 min around each time point of a trajectory. Trajectories shorter than 10 min, and the first and last 5 min of each trajectory (which are prone to noise), were thus not included. The experimental time axis was divided into 20 min intervals and the mean GR for each bacterium within each time interval was calculated. The distribution of GRs and Lengths across the population within each time interval was shown in boxplots.

*GR(t)*, and length, *L(t)*, were fitted by Gompertz functions, the functional dependencies of which from the duplication time was estimated through *T*_2_(*t*) = (2⁄3) × (*L_final_*/*GR*(*t*)). The number of bacteria on the surface at any time after binding, *N(t)*, will depend both on *T_2_(t),* and on the rate of bacteria release from the surface, indicated by the mean binding time, *T_½_*, through the relation

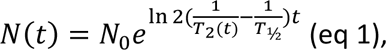

where *N_0_* is the number of bacteria at the surface right before bacteria starts to divide, which was approximated by the time of the inflection point of the Gompertz fit. *T_½_* was determined from the best fit of eq. 1 to plots of the number of bacteria versus time.

## Supplementary Information

Supplementary Figure 1: ^1^H NMR spectrum of AMC-25-04 in DMSO-*d6*. Supplementary Figure 2: Calculated flow profiles in the microfluidic channel Supplementary Figure 3: Bacterial alignment and small movement for *E. coli-Fim+* Supplementary Figure 4: Growth rates for *WT E. coli* in LB + 100 µM AMC-2504

Supplementary Movie 1: Small motions of *WT E. coli* bound to mannose-modified surface (25 times real speed)

Supplementary Movie 2: Shrinking bacterium (250 times real speed)

Supplementary Movie 3: Small motions of *WT E. coli* bound to AMP-modified surface at 30 min (25 times real speed)

Supplementary Movie 4: Small motions of *E. coli-ΔFimA* bound to AMP-modified surface at 30 min (25 times real speed)

Supplementary Movie 5: Small motions of *E. coli-ΔFimA* bound to AMP-modified surface at 160 min (25 times real speed)

Supplementary Movie 6: Small motions of *WT E. coli* bound to AMP-modified surface at 160 min (25 times real speed)

## Acknowledgements

Marta Tous Mohedano and Dr. Anne Farewell are acknowledged for constructing bacterial strain MG1655ΔfimA. The authors are grateful for analytical assistance from RISE scientist Dr. P. Borchardt. RISE scientist Dr. Per Borchardt is acknowledged for assistance with analysis of ToF-SIMS data. This research was financed by the Swedish Research Council (Grant 2019-05215).

## Author contribution statement

AH, MB & AL planned and implemented the live-microscopy experiments and the analysis. EAK, WS & JSS planned and implemented the synthesis of the antimicrobial peptides. All authors took part in writing the manuscript.

