## Supplemental Figures for "Preventing *E. coli* biofilm formation with antimicrobial peptide surface coatings: recognizing the dependence on the bacterial binding mode using live-cell microscopy"

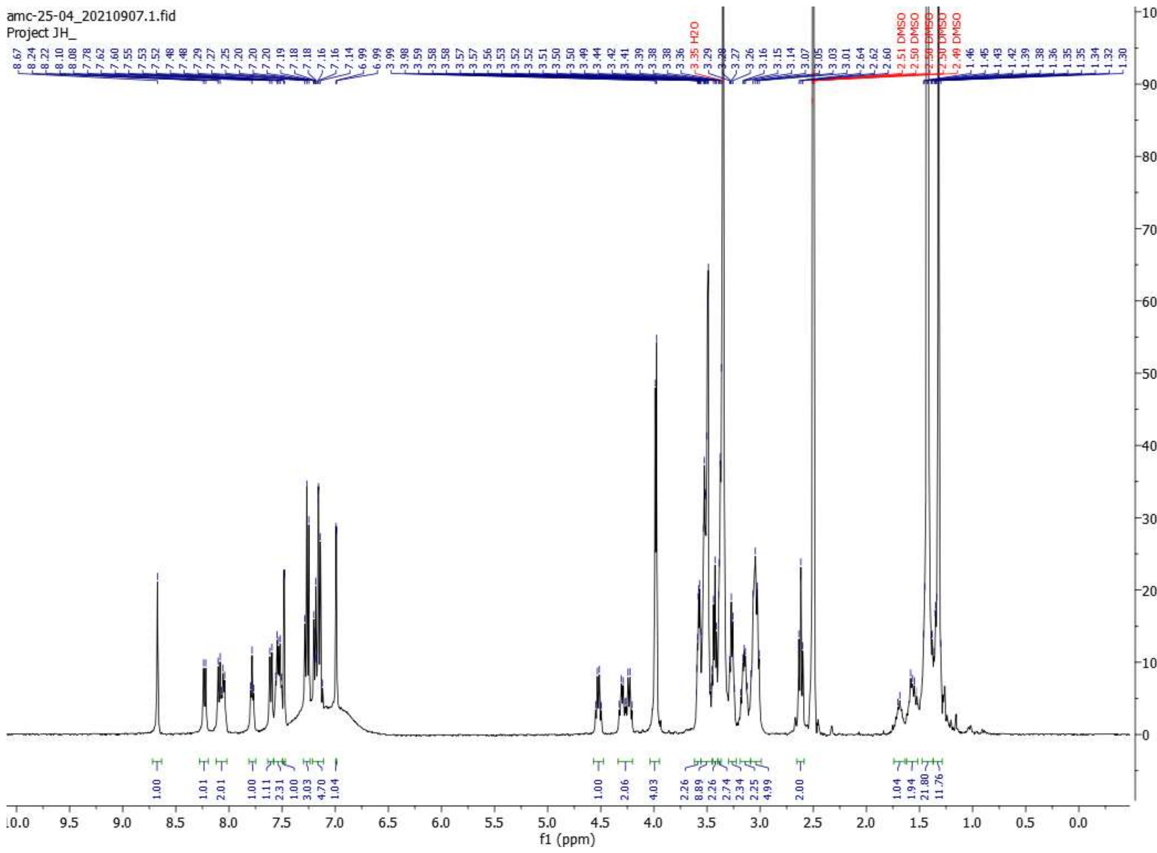

Supplementary Figure 1: $^{1}H$ NMR spectrum of AMC-25-04 in DMSO-*d6*.

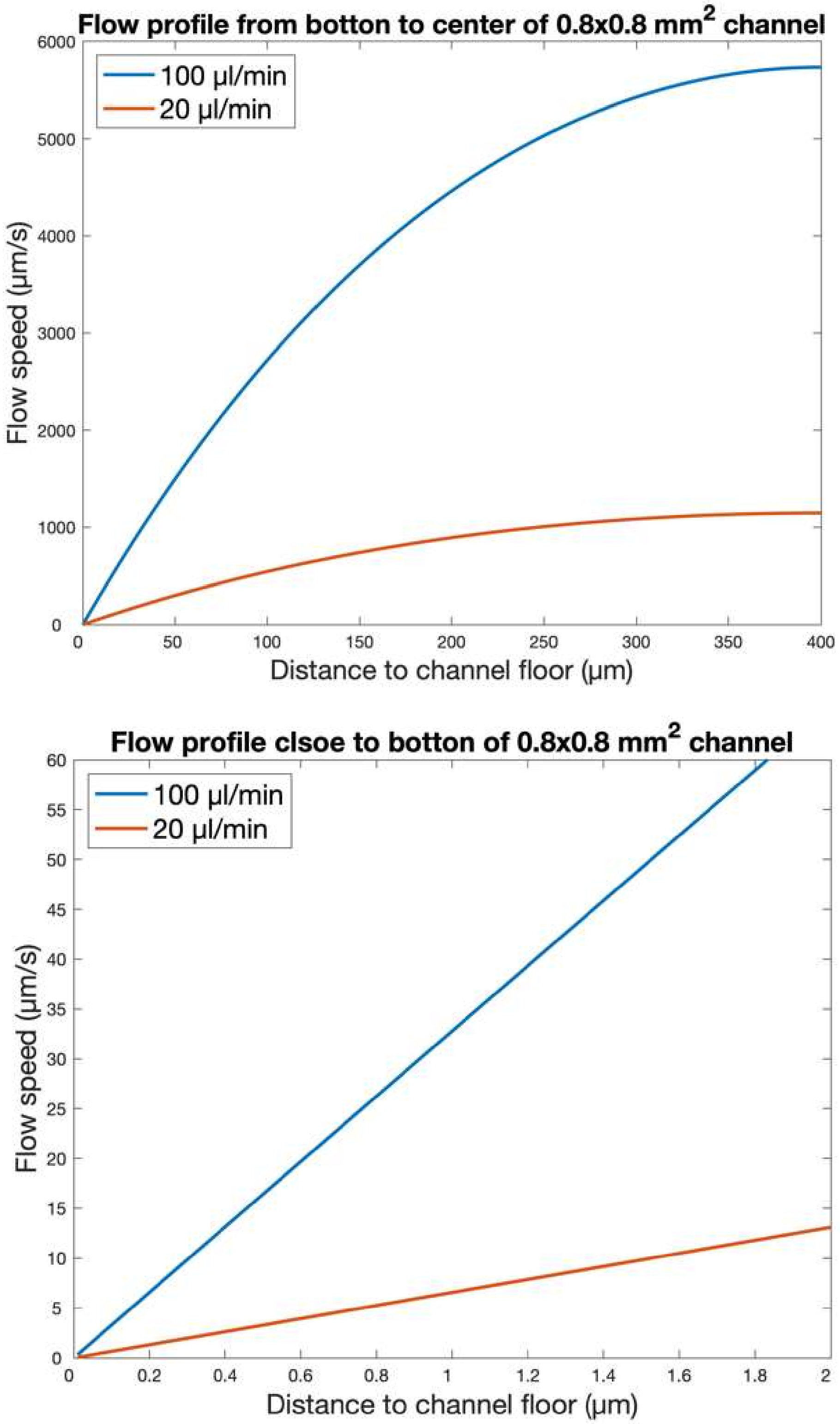


Supplementary Figure 2: Calculated flow profiles showing the flow speed profile in the bottom half of the channel (upper panel) and close to the bottom of the channel where the bacteria stick (lower panel).

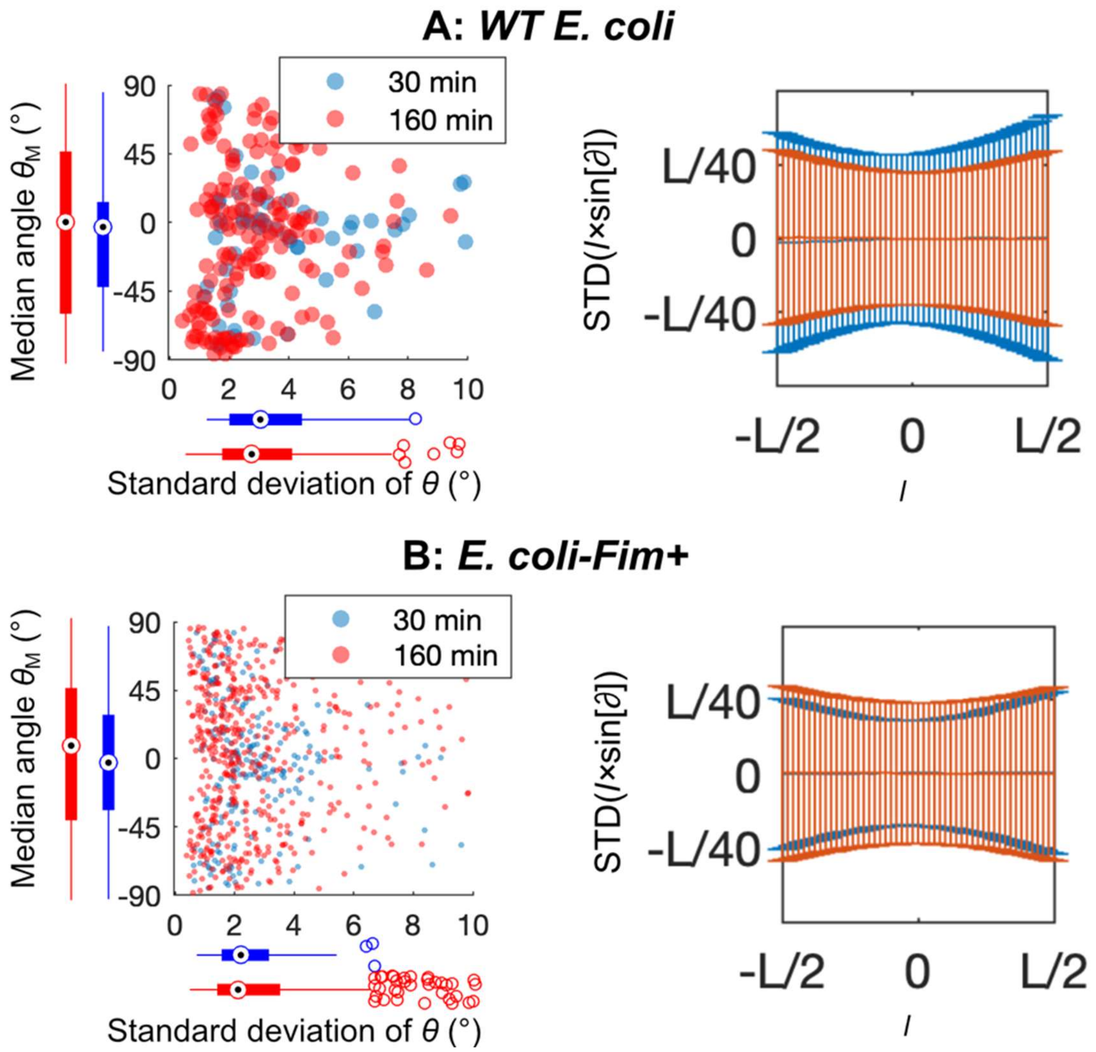


Supplementary Figure 3: The combined scatter and boxplots show the median angle, $\theta_M$, versus the standard deviation (STD) of $\theta$ for all bacteria at an early (30 min, blue points) and a late (160 min, red points) timepoint of the experiment. The bar plots to the right show the distribution (standard deviations) of the instantaneous separations, $l \times \sin(\partial\theta)$, for all positions l of all bacteria present early (30 min, blue bars) and late (160 min, red bars) in the experiment. **(A)** *WT E. coli* **(B)** *E. coli-Fim+*.

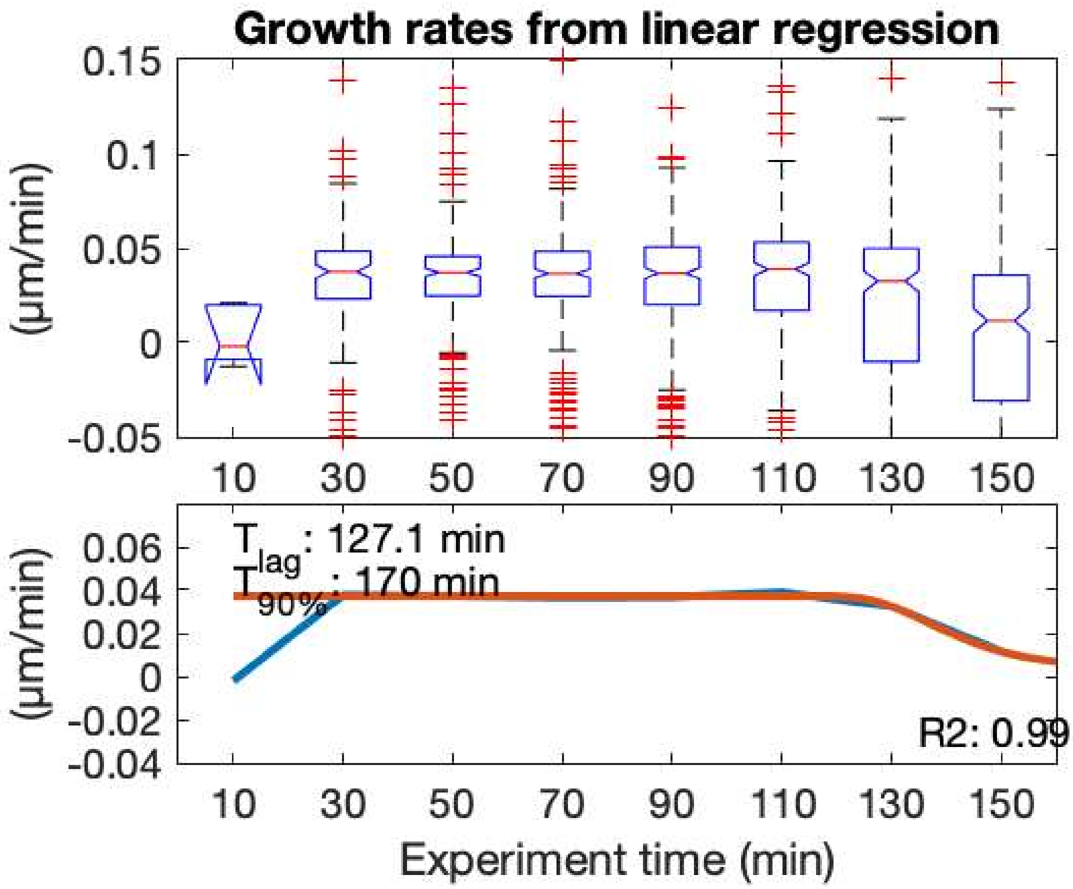


Supplementary Figure 4: Boxplot showing the distribution of GRs of the surface-bound bacteria charged with 100 µM AMC-2504 in the LB growth media. Data for t<20 min. i.e. during the injection phase, is not representative since few bacteria could be traced >10 min during the binding phase. Note that the distribution of GRs is narrow in the beginning of the experiment but broadens towards the end.
